# Dissociating ego- and allocentric neglect after stroke: prevalence, laterality and outcome predictors

**DOI:** 10.1101/200386

**Authors:** Nele Demeyere, Celine R. Gillebert

## Abstract

Visuospatial neglect is a neuropsychological condition commonly experienced after stroke, whereby a patient is unable to attend to stimuli on their contralesional side. We aimed to investigate whether egocentric and allocentric neglect are functionally dissociable and differ in prevalence, laterality and outcome predictors. A consecutive sample of 366 acute stroke patients completed the Broken Hearts test from the Oxford Cognitive Screen. A subsample of 160 patients was followed up 6 months later. We evaluated the association between egocentric and allocentric neglect, contrasted the prevalence and severity of left-sided versus right-sided neglect, and determined the predictors of persistence versus recovery at follow up. Clinically, we found a double dissociation between ego- and allocentric neglect, with 50% of the neglect patients showing ‘only’ egocentric neglect and 25% ‘only’ allocentric neglect. Importantly, patients with only allocentric neglect did not demonstrate any egocentric spatial bias in the locations of the allocentric errors. Left-sided egocentric neglect was more prevalent and more severe than right-sided egocentric neglect, though right-sided neglect was still highly prevalent in the acute stroke sample (35%). Leftsided allocentric neglect was more severe but not more prevalent than right-sided allocentric neglect. Overall recovery of neglect was high: 81% of egocentric and 75% of allocentric neglect patients recovered. Severity of neglect was the only significant behavioural measure for predicting recovery at 6 months.

## Introduction

Visuospatial neglect is highly prevalent after stroke. The condition is associated with an attentional deficit where patients fail to orient, perceive and interact with stimuli on the contralesional side of space. A systematic review of 17 studies comparing the prevalence of neglect acutely, after left (LBD) and right brain damage (RBD), found that the median prevalence of left-sided neglect (after RBD) was twice as high as for right-sided neglect (43% vs 21%) [1]. A recent study on 335 subacute patients (median 28 days post stroke) found incidences of 9% right neglect and 16% left neglect [2], though prevalence reports for right neglect have varied from 2% [3] to 65% [4] after LBD. Bowen and colleagues suggested potential inclusion criteria and selection biases may explain the variability and noted the problematic small samples. Suchan, Rorden and Karnath [5] specifically studied 48 patients with focal LBD and set out to include patients with aphasia in their acute sample. They found a right neglect prevalence of 44% and no statistical difference in the severity of neglect between LBD and RBD patients [6].

Neglect appears to be a heterogeneous syndrome with dissociable symptoms. One such distinction is between space-based (ego-centric) and object-based (allo-centric) neglect. Egocentric neglect is where the patient fails to attend to the contralesional side of space (with reference to their own body midline). Allocentric neglect is where the patient fails to attend to the contralesional side of an object in focus. Double dissociations have been reported between patients and even within a single bilateral patient who demonstrated egocentric neglect on one side and allocentric neglect on the other side [7]. Work by Kleinman and colleagues [8] suggested that, although RBD results mainly in left egocentric neglect, LBD more commonly resulted in right allocentric neglect. This may even help explain why the prevalence of right neglect has been underestimated, as allocentric neglect is often not explicitly assessed [see overview by 9]. In addition, neuroanatomical lesion studies have supported the dissociation [e.g. 10]. A recent meta-analysis of 22 lesion-symptom mapping studies concluded that patients with more anterior lesions experience egocentric neglect, whereas patients with more posterior lesions experience allocentric neglect [11]^1^.

Taking an alternative view, ego- and allocentric neglect may reflect two aspects of a central underlying disorder [12], where allocentric neglect is simply occurring when the attentional window is narrowed to a single object [13] or where allocentric biases are modulated by their egocentric position [14,15]. Relatively strong correlations between ego-and allocentric neglect have been reported [16] and it has been argued that the two cannot be fully dissociable given findings where the allocentric deficit was worse for stimuli in the contralesional compared to the ipsilesional side, supporting the notion that allocentric biases occur due to a spatial gradient of attentional weights (see Figure 1).

**Figure 1.**
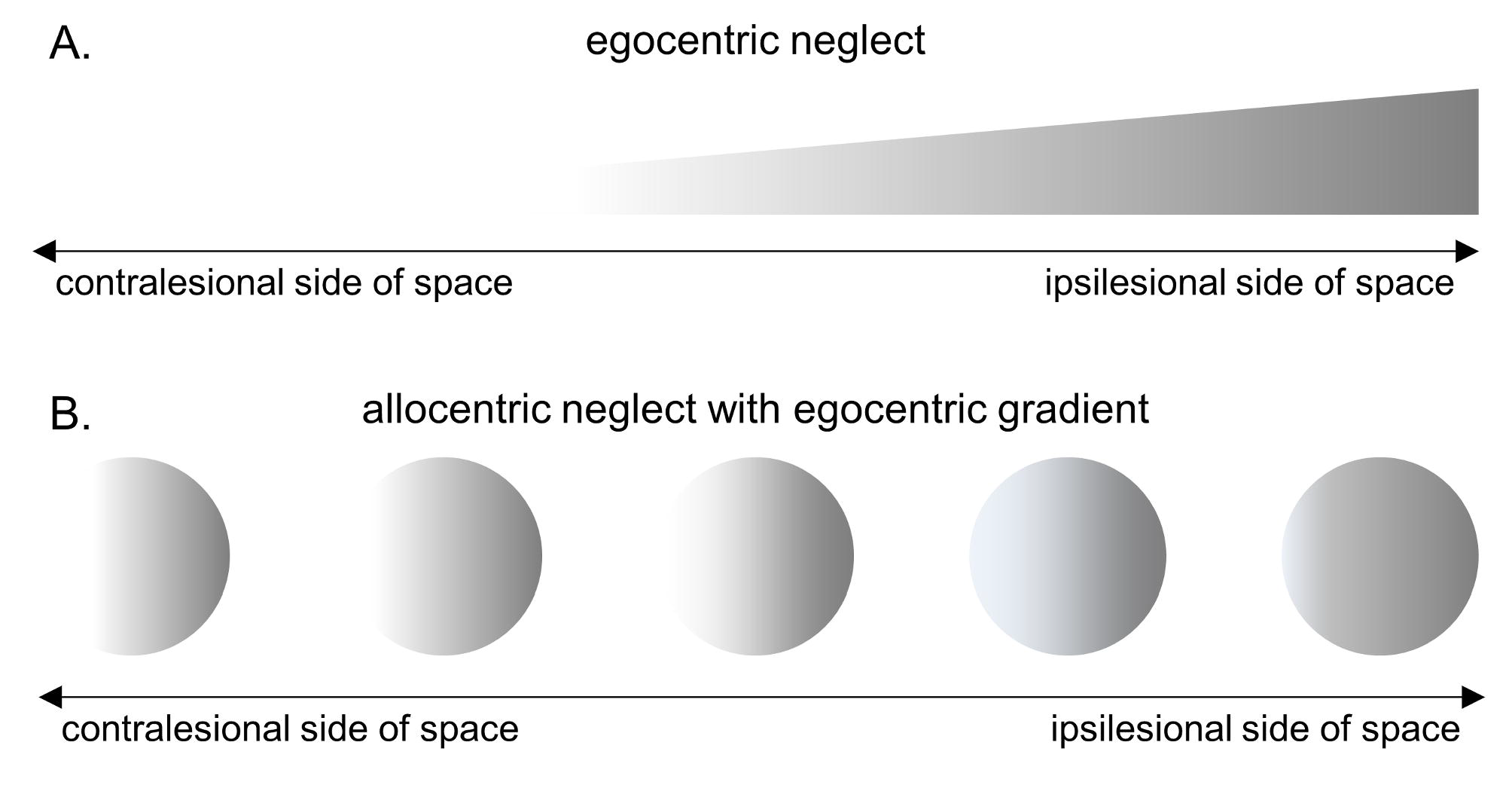
Attentional gradient. Illustration of how an egocentric spatial gradient (A) could account for allocentric errors (B).

In this study, we aimed to contrast the prevalence and severity of left- and right-sided egocentric and allocentric neglect in a large sample of acute stroke patients. In addition, we investigated the distributions of allocentric errors in egocentric space, and assessed recovery rates and behavioural predictors of persistent neglect at 6 months.

## Material and Methods

### Participants and procedure

This study was designed as a cross-sectional observational study in 419 stroke patients recruited from two acute stroke units at the John Radcliffe Hospital Oxford and the University Hospital Coventry and Warwickshire between 2012 and 2014. Patients completed the Oxford Cognitive Screen (OCS) [17] on the acute ward and agreed to be contacted 6 months later for a follow-up visit. Patients were recruited consecutively, depending on practical availability of the patient and researcher at the time. The research team was present for several hours every weekday to recruit the patients, they were under clear instructions to try to see everyone that was able to be seen, thereby avoiding a selection bias based on behavioural symptoms or type of stroke. Though we collected the clinical admission CT scans for the sample, only a subset proved to be of high enough quality to determine the more precise lesion location. We chose not to restrict the analysis to these patients, as the purpose of this study was to assess the prevalence, laterality and recovery of neglect in an unbiased sample reflecting the clinical reality [see also 18]. This is in contrast with studies investigating the precise relationship between neuropsychological symptoms and lesion location who typically exclude patients with a prior stroke, multifocal lesions or excessive lacunae, tiny lesions, medical comorbidities or contraindications for MRI [e.g. 19,20].

Inclusion criteria for the study were: patients should be within 3 weeks of a confirmed diagnosis by clinicians to have had an ischaemic or haemorrhagic stroke, be able to concentrate for 15 minutes and be able to give written informed consent themselves (or witnessed consent in cases of motor problems or agraphia). 366 patients of the consented sample were able to complete the Broken Hearts Test of the OCS. Demographics data and stroke information were collected from the medical notes (Table 1). Reasons for exclusion are given in Supplementary Table 1.

**Table 1:** Demographics and stroke data

|  | acute<br>(N=366) | follow-up<br>(N=160) |
| --- | --- | --- |
| Gender (male \| female) | 192\|174 | 91\|69 |
| Age (years: mean ± SD) | 73±14 | 72±13 |
| Stroke to test interval (days: mean ± SD) | 6±4 | 193±49 |
| Lesion side (left \| right \| bilateral \| not specified) | 148\|181\|24\|13 | 61\|79\|24\|9 |

Six months later, all patients were contacted by telephone and verbal consent was taken before arranging a homevisit with a research assistant to complete follow up cognitive assessments, among which the Apples Cancellation Task of the Birmingham Cognitive Screen (BCoS) [21]. 160 patients completed the follow up (see Supplementary Table 2 for drop out reasons).

This study was approved by the National Research Ethics Service (Ref: 11/WM/0299; Protocol RP-DG-0610-10/046).

### Cancellation task

The Broken Hearts Test [17] and Apple Cancellation Test [21] are equivalent, highly sensitive cancellation tests assessing ego- and allocentric neglect. The task is to strike through the complete shape outlines (n=50) amongst distractor shapes with gaps on the right (n=50), or the left (n=50) of the contour (Supplementary Figure 1). The items are positioned semi-randomly on an A4 landscape page, equally distributed over a virtual grid. Patients are given up to two practices with demonstration and thorough explanation to ensure they understood the task before starting the test. There is a time limit of three minutes in which the task was to be completed.

We calculated the following outcome measures (Figure 2)

**Figure 2.**
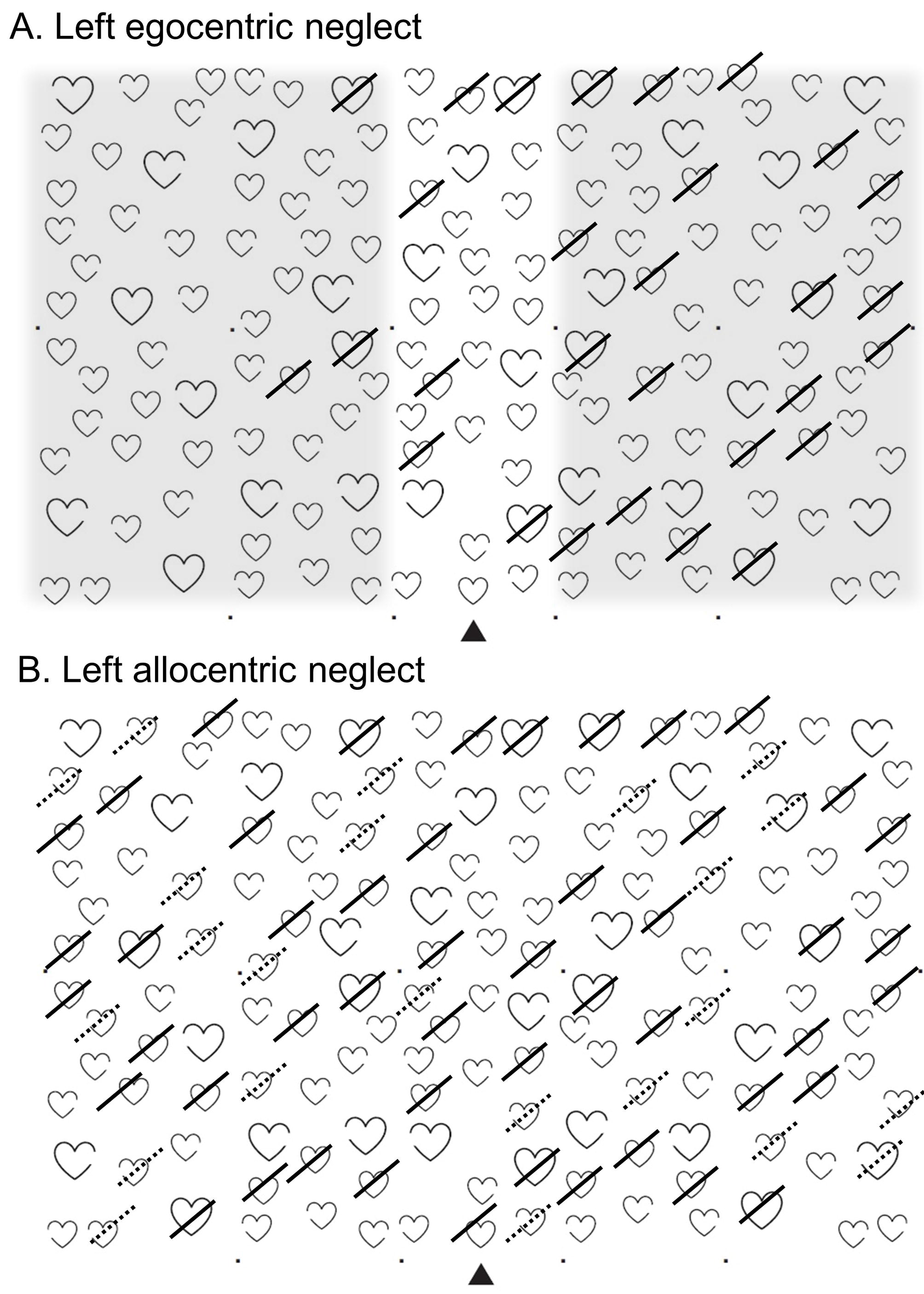
The Broken Hearts Test. (A) Egocentric neglect is operationalized as an asymmetry value calculated by subtracting the number of hits (full strikes) on the left page side from the hits on the right hand side, also taking into account the total number of hits (overall correct should be lower than 42, based on cut off from normative data). Only the shaded areas are taking into account to calculate the egocentric asymmetry score. Note that the shaded areas have only been added to clarify the scoring; the page presented to the patients contains only the hearts. (B) Allocentric neglect is operationalised as an asymmetry score calculated by subtracting the number of false positives (dashed strikes) with a left gap from the number of false positives with a right gap.

1. **Hits:** The number of correctly cancelled full outlines (hearts or apples) (max 50);
2. **Allocentric errors:** The number of incorrectly cancelled distracters (max 100);
3. **Egocentric asymmetry:** The difference between the number of hits on the left versus right side of the page. Only the shaded areas in Figure 2A were taken into account. Positive values denote left-sided neglect, negative values right-sided neglect;
4. **Allocentric asymmetry:** The difference between the number of allocentric errors with a left versus right gap. Positive values denote left-sided neglect, negative values right-sided neglect;
5. **Egocentric asymmetry of allocentric errors:** The difference between the number of allocentric errors on the left versus right side of the page.

Cut-offs for impairment were derived from 5^th^/95^th^ centile scores from the normative data (see 17). Patients were considered to have ‘egocentric neglect’ if they had an accuracy score < 42 *and* an absolute egocentric asymmetry score ≥3. Patients were considered to have ‘allocentric neglect’ with an absolute allocentric asymmetry ≥2^2^.

## Results

### Dissociable subtypes of egocentric and allocentric neglect in acute stroke

48% of the patients who completed the Broken Hearts Test, demonstrated neglect (Figure 3A). Half of them presented with only egocentric neglect, one quarter with only allocentric neglect and a further quarter with both egocentric and allocentric neglect.

**Figure 3.**
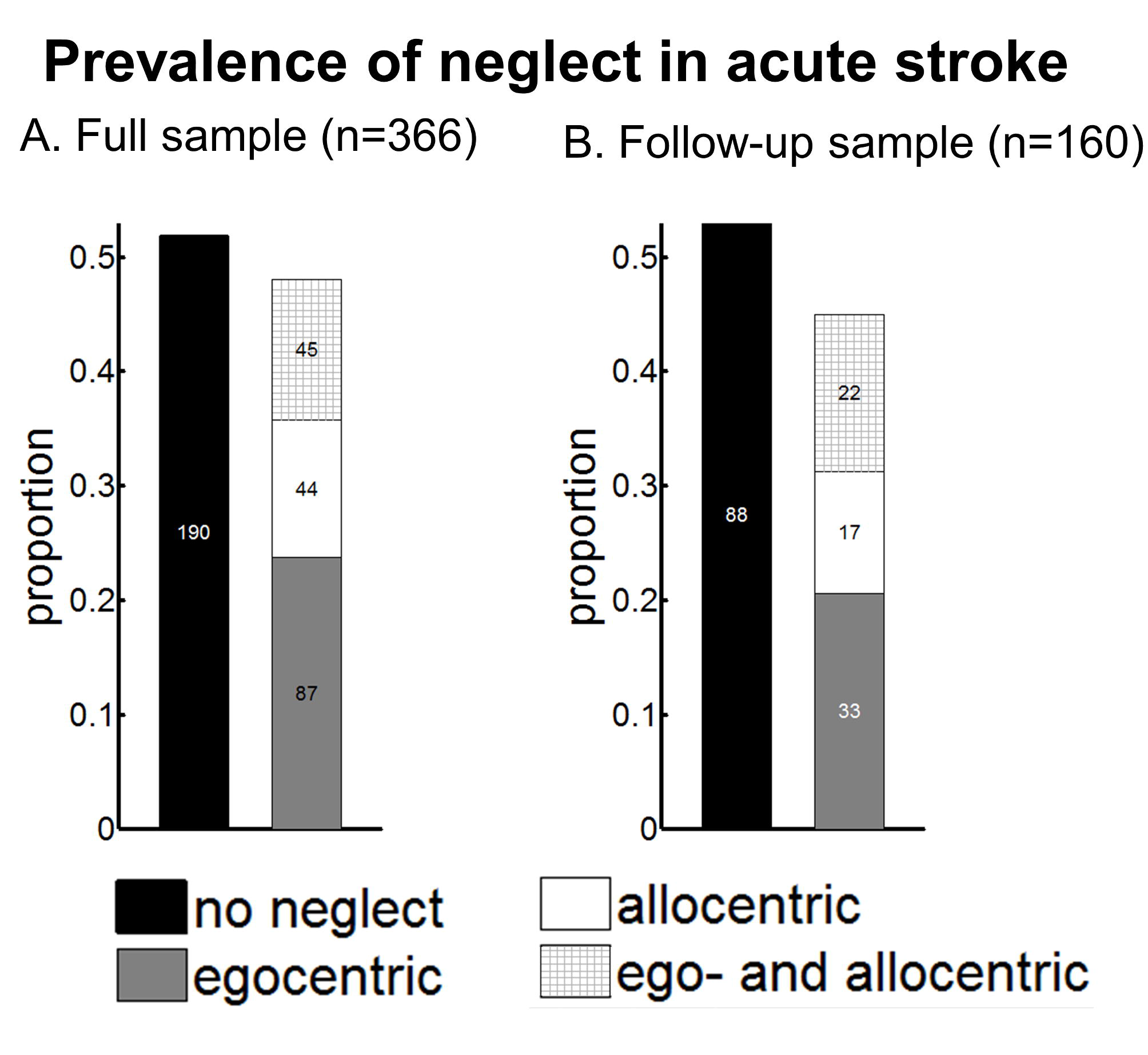
Prevalence of ego- and allocentric neglect in a consecutive sample of 366 acute stroke survivors (A) and a subsamble of 160 acute stroke survivors who received a follow-up at 6 months (B).

To assess the relationship between the severity of ego- and allocentric spatial biases in a more sensitive way, we correlated the ego- and allocentric asymmetry scores in the three groups (Figure 4). The patients with both ego- and allocentric neglect demonstrated a clear correlation between the asymmetry scores (N=45, r=.55, p<0.0001), however no correlation was present in patients with only egocentric or only allocentric neglect (N=131, r=.02, *p*=0.78).

**Figure 4.**
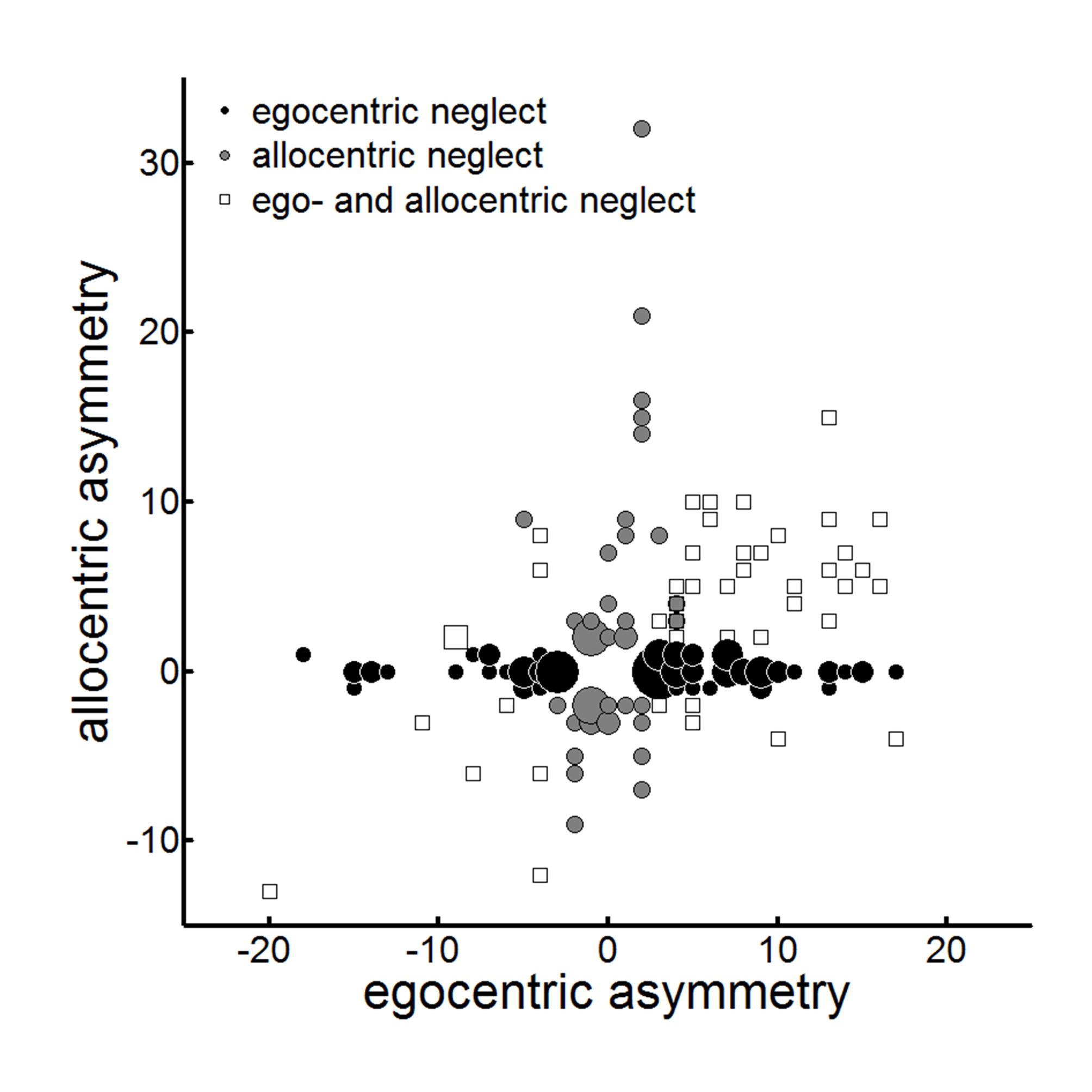
Egoversus allocentric neglect. Bubble plot of egocentric versus allocentric asymmetry scores in the 176 patients with neglect. Positive values reflect left-sided neglect, negative values reflect right-sided neglect. Legend: black circle: patients with only egocentric neglect (N=87); grey circle, patients with only allocentric neglect (N=44); white square: patients with ego- and allocentric neglect (N=45).

To determine whether the false positives made by patients with allocentric neglect were asymmetrically distributed in egocentric space, we examined where on the page the allocentric errors were made. An ANOVA with side of space (same vs opposite side of allocentric errors) and presence of egocentric neglect (allocentric only, ego- and allocentric) showed a main effect of side of space (F(1,76)=34.91, p<0.001) and an interaction between the presence of egocentric neglect and side of space (F(1,76)=43.83, p<0.001) but no main effect of group (F(1,76)=0.34, p=.56) (Figure 5). Post-hoc t-tests suggested that for the allocentric only group, the average number of errors on each side of the page was the same (t(42)=0.70, p=0.49). This demonstrates that no egocentric bias was present to explain the allocentric asymmetry, in contrast to the theoretical framework outlined in Figure 1. In contrast, patients with both ego- and allocentric neglect made fewer errors on the egocentrically neglected side of the page (t(34)=-6.82, *p*<0.001).

**Figure 5.**
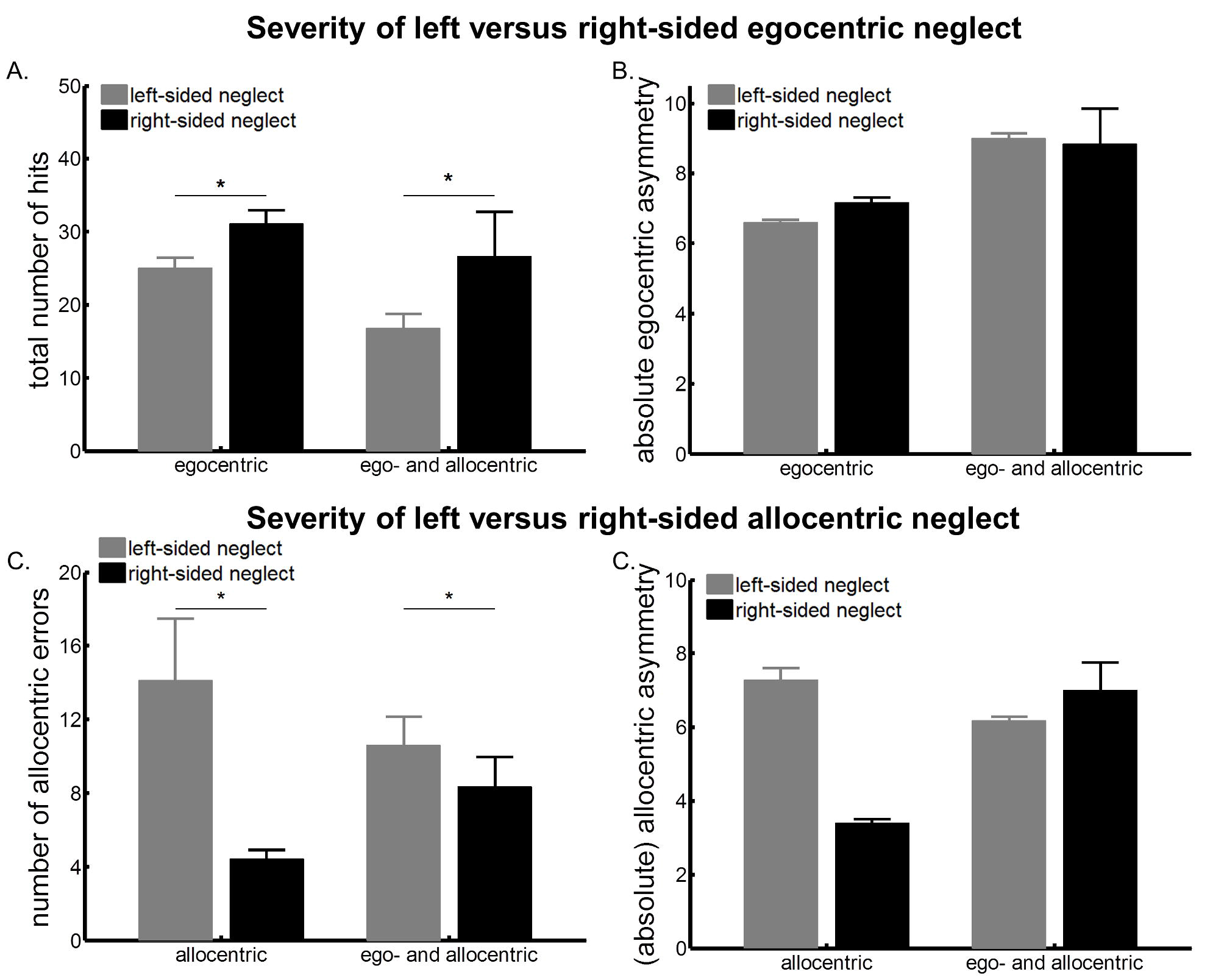
Egocentric bias in the allocentric errors. Number of allocentric errors according to the side of space (same versus opposite side of the allocentric errors) for patients with allocentric neglect only and patients with both ego-and allocentric neglect.

### Prevalence of left-sided and right-sided neglect

In patients with only egocentric neglect, the prevalence of right-sided neglect (35%) was significantly lower than that of left-sided neglect (65%) (χ^*2*^(1)=8.38, *p*<0.004). Similar results were obtained in patients with ego- and allocentric neglect: 74% presented with left-sided and 26% presented with right-sided neglect (χ^*2*^(1)=15.11, *p[*0.001). In contrast, there was no difference in the prevalence of left-sided (56%) and right-sided (44%) neglect in patients with only allocentric neglect (χ^*2*^(1)=.36, *p*=0.55).

### Severity of left-sided and right-sided neglect

To directly compare the severity of left- versus right-sided egocentric neglect, we submitted the number of hits and the egocentric asymmetry score to an ANOVA with neglected side (left, right) and the presence of allocentric neglect (egocentric only, ego- and allocentric) as factors.

We observed a main effect of neglected side on the number of hits (F(1,118)=8.63, *p*=0.004) but not on the egocentric asymmetry score (F(1,118)=0.04, p=.85). Patients with right egocentric neglect demonstrated a higher overall accuracy than patients with left egocentric neglect, but the strength of the spatial bias did not differ between the groups (Figure 6A-B). In addition, we observed a main effect of group (hits: F(1,118)=5.43, p=.02; asymmetry: F(1,118)=3.76, *p*=.06), showing that patients with ego- and allocentric neglect performed worse on the cancellation task than patients with only egocentric neglect. The interaction between both factors was not significant (hits: F(1,118)=.48, *p*=.49; asymmetry: F(1,118)=0.12, *p*=0.73), suggesting that having added allocentric on top of egocentric neglect does not lead to a worse egocentric cancellation performance than having egocentric neglect only.

**Figure 6.**
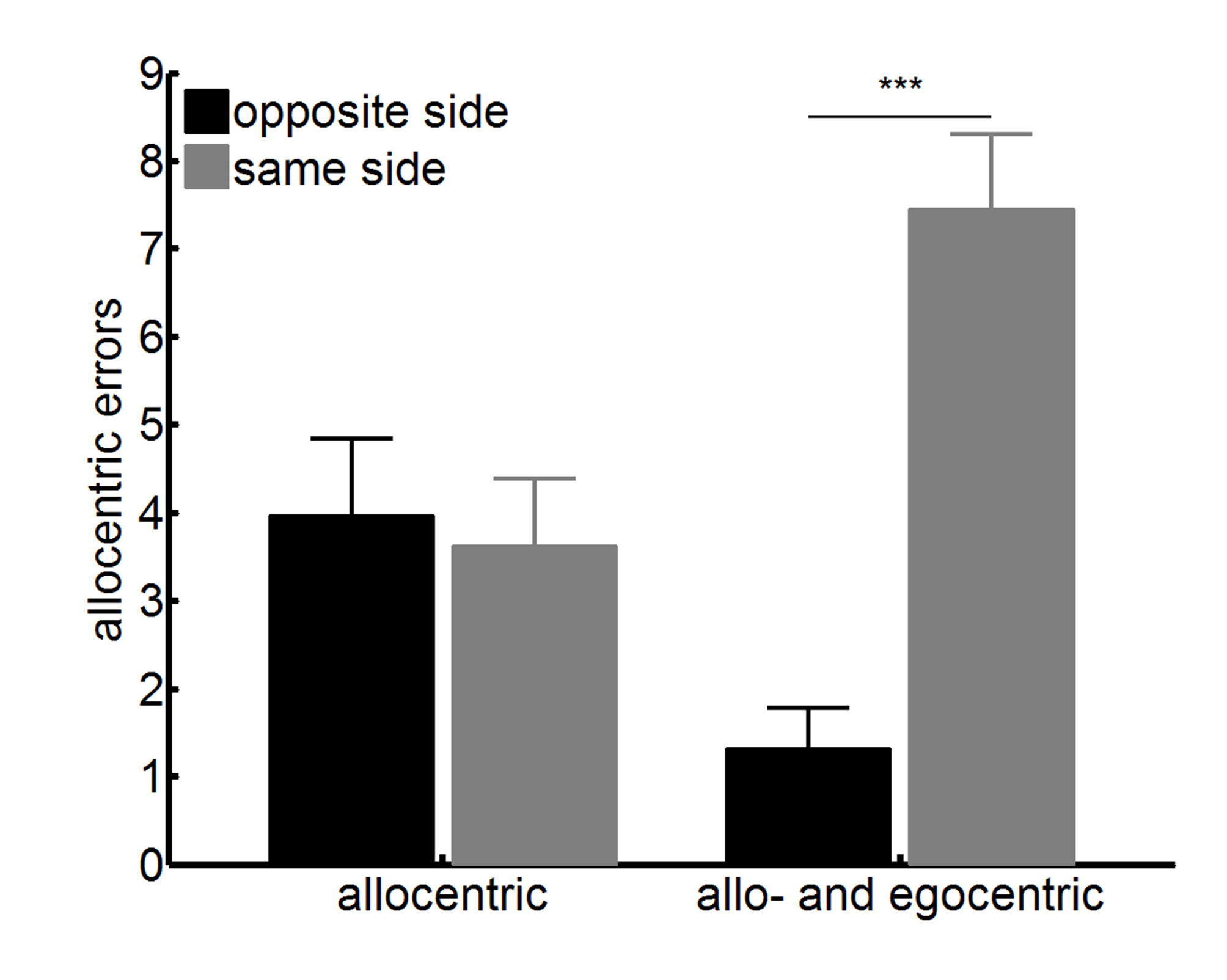
Left versus right-sided neglect. (A-B) Severity of egocentric neglect is reflected in the total number of hits (A) and the absolute egocentric asymmetry score (B). (C-D) Severity of allocentric neglect is reflected in the total number of allocentric errors (C) and the absolute allocentric asymmetry score (D).

To contrast the severity of leftversus right-sided allocentric neglect, we submitted the number of allocentric errors and the allocentric asymmetry score to an ANOVA with neglected side (left, right) and the presence of egocentric neglect (allocentric neglect only, ego- and allocentric) as factors.

There was a significant main effect of neglected side on the number of allocentric errors (F(1,75)=3.30, *p*=.039), but not on the allocentric asymmetry score (F(1,75) = 1.39, *p=0.*24) (Figure 6C-D). The presence of egocentric neglect did not affect the allocentric outcome variables (errors: F(1,75)=0.005, *p*=.95; asymmetry: F(1,75)=0.91, *p* = .34), and did not interact with the neglected side (errors: F(1,75)=1.71, *p*=0.20; asymmetry: F(1,75)=3.29, *p*=0.07).

To summarize, our data demonstrated a higher prevalence of left compared to right egocentric neglect, but a similar prevalence of left compared to right allocentric neglect in this acute sample. The severity of the neglect was more prominent in leftsided neglect (lower accuracy and/or more allocentric errors).

### Recovery of ego- and allocentric neglect

For the patients who completed the follow-up assessment 6 months later, the distribution of acute prevalence rates of the different neglect types was proportionally the same as in the full sample of patients (Figure 3B). 55 of these patients demonstrated egocentric neglect at the acute stage, and only 11 were still impaired at follow-up. Allocentric neglect was present in 39 patients acutely and remained present in 10 patients at follow-up. This demonstrates very high recovery rates, at 81% and 74% for egocentric and allocentric neglect respectively.

To predict recovery from ego- or allocentric neglect from the initial behavioural profile, we carried out a binary logistic regression. To predict the recovery of patients who suffered from egocentric neglect in the acute phase of stroke, five factors (age, hits, absolute egocentric asymmetry, neglected side, presence of allocentric neglect) were entered into the regression logarithm. Only the number of hits in the acute phase of stroke was a significant predictor of outcome 6 months later (*p*=0.007), with a trend present for age as well (*p*=0.06). Persistent neglect was associated with lower performance and higher age in the acute phase after stroke. Noteworthy, an ANOVA on the number of hits with time-point (acute, follow up) and persistence (recovered from neglect, persistent neglect) as factors revealed a main effect of time-point (F(1,53)=52.90, *p*<0.001) and persistence (F(1,53)=29.96, *p*<0.001) but no interaction between time-point and persistence (F(1,53)=0.90, *p*=0.35). In other words, patients with persisting neglect did show a similar degree of improvement compared to patients who recovered from neglect (Figure 7A).

**Figure 7.**
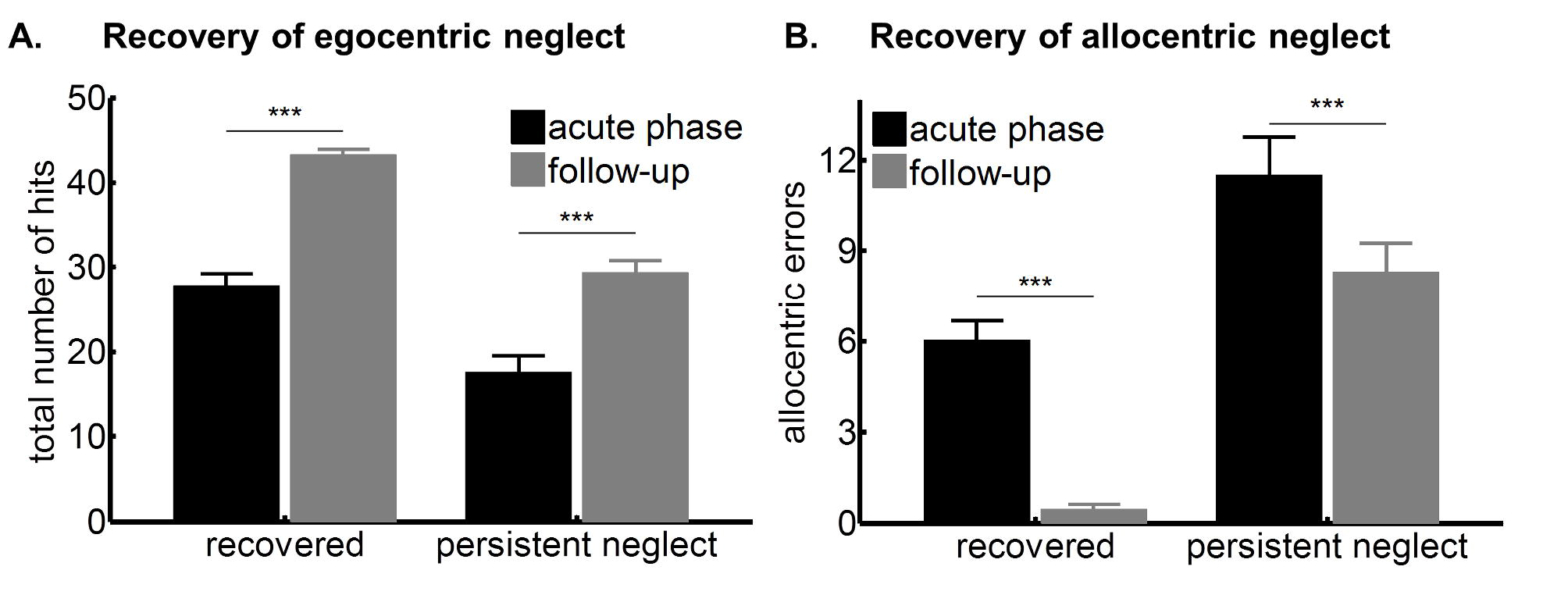
Recovery of ego- and allocentric neglect. (A) Total number of hits in the acute phase after stroke and at follow-up for patients who recovered from egocentric neglect and in patients with persistent neglect. (B) Number of allocentric errors in the acute phase after stroke and at follow-up for patients who recovered from allocentric neglect and in patients with persistent neglect.

Similarly, for allocentric neglect, four factors (age, allocentric errors, neglected side, presence of egocentric neglect) were entered into the regression logarithm. The allocentric asymmetry score was not included as it highly correlated with the number of allocentric errors (r=.77, p<.001). Only the number of allocentric errors significantly predicted the persistence of allocentric neglect at follow up (*p =* .02). Given that left-sided allocentric neglect was associated with more allocentric errors, this particular combination had the worst prognostics: 10 out of 26 patients remained impaired at follow-up. An ANOVA on the number of allocentric errors with time-point (acute, follow-up) and persistence (persistent neglect, recovered from neglect) as factors showed a main effect of time-point (F(1,37) = 19.02, *p*<0.001) and persistence (F(1,37)=24.73, *p*<0.001) but no interaction between time point and persistence (F(1,37)=1.40, *p*=0.24). As can be seen in Figure 7B, patients with persistent neglect also improved substantially between the acute and the chronic phase.

In summary, our analyses suggest that the overall accuracy on the cancellation task (in terms of the number of hits and allocentric errors) was the only significant predictor of recovery at 6 months.

## Discussion

This study set out to address a series of important questions relating to the nature of hemi-spatial neglect, its prevalence and its outcome predictors. We included a large sample of acute stroke patients who were not selected based on lesion location or behavioural profile and followed up a subset of them 6 months later. This is the biggest sample study to assess acute left and right neglect prevalence, severity and recovery to date.

We found clear evidence to suggest that egocentric and allocentric neglect are subserved by separate underlying processes. First, we documented a double dissociation, with substantive groups of patients with ‘only’ egocentric and ‘only’ allocentric neglect, where there was no relation between asymmetry values for ego- and allocentric asymmetries. This finding may seem in contrast to the overall correlation (r=0.35) between ego-and allocentric reported by Rorden and colleagues [16]. We note however that their sample included only 32 RBD patients. It may be that this smaller sample accidentally contained more patients with both types of neglect. We indeed also observed a clear correlation in the sub-group presenting with both ego- and allocentric neglect (r=.55). We further demonstrated that in patients with allocentric neglect only, there was no spatial asymmetry to the distribution of the errors on the page. Where a previous study [14] found an amelioration of allocentric neglect symptoms at more ipsilesional egocentric positions, the 3 patients in that study presented with both types of neglect. Here, in the allocentric only patients we did not find any evidence for an egocentric exploration bias. This contradicts the theory proposed by Li and colleagues [15] that allocentric neglect appears due to an overall spatial gradient (Figure 1). Instead we suggest that our findings provide strong behavioural evidence for truly dissociable neglect types. The evidence is in line with previous smaller studies [22] and a striking dissociation in a single case study[23]. These behavioural dissociations complement the neuro-anatomical findings for separable underlying mechanisms [24].

The allocentric neglect observed here may reflect an object-based neglect or spatial neglect within a smaller reference frame [13]. Allocentric neglect can indeed be considered in terms of local and global visual representations. For instance, impaired detection of the gap in the Broken Hearts and Apples Cancellation Tasks may stem from impaired attention to one side of the local spatial representation [25]. In this case, ego- and allocentric neglect may share a common spatial deficit (which could be viewed as the core symptom of neglect [12]). The heterogeneity in neglect in the level of representation is then due to the manifestation of this core deficit, either at a global or local scale, or an egocentric or allocentric viewpoint. Even when the dissociation observed here would be viewed as reflecting a difference in ‘satellite symptoms’ of neglect, with a common core spatial deficit, the experience of the two results in different functional impairments: missing half of an object anywhere in space is existentially different from a global failure to attend to one side of space and is likely to require a different rehabilitation protocol.

In contrast to most studies on neglect, we did not restrict inclusion to the study to patients with only RBD. Interestingly, this unbiased sample in acute stroke revealed high incidences of right-sided ego- and allocentric neglect. Nevertheless, it was clear that left-sided neglect symptoms were more severe in terms of a lower number of hits and a higher number of allocentric errors.

These findings are in line with long standing theories of a right hemisphere dominance of hemispatial neglect. Our findings are comparable to those by Stone et al [4], Baldassare and colleagues [26], and Ten Brink et al [2] who found a considerable number of patients with right-sided neglect in the acute stage, though their symptoms were less severe. Particularly in the chronic stage the frequency and severity of left-sided neglect is notably higher. This hemispheric difference was initially explained by interhemispheric competition after RBD leading to an imbalance in attentional control, where the right side of space is ‘over’ attended [27]. A different account was proposed by Mesulam [28] in terms of spatial coding for the left side of space being subserved only by the right hemisphere, whereas spatial coding for the right side of space is subserved by both hemispheres. Consequently, LBD means the right hemisphere can take over the spatial representations. A different account of the right lateralization is that the neglect syndrome encompasses more than a spatial attention bias alone. The presence of impaired sustained attention and working memory in patients with neglect [29], processes that are right lateralized, may contribute to the higher severity and the worse recovery of left-sided as opposed to right-sided neglect [30]. Several studies have now shown the presence of spatial asymmetries in LBD patients without neglect (e.g. see [31,32]).

We found high rates of recovery for ego- and allocentric neglect. Importantly, we also observed substantial improvement in the subset of patients with persisting neglect. When predicting the likelihood of recovery based on the behavioural profile in the acute phase after stroke, only the symptom severity on the initial measures was predictive. Patients with more severe neglect symptoms were less likely to recover. Our observations may have significant implications for research into neglect rehabilitation. Given that the majority of neglect patients are likely to recover, rehabilitation may either be more useful in a later stage, or acute rehabilitation of neglect should target those with the more severe neglect symptoms as they have the worst prognosis for recovery.

A second point relates to left allocentric neglect specifically, which was most likely to be persistent. All right allocentric patients had recovered, but 48.5% of left allocentric neglect was persistent 6 months later. Bickerton and colleagues [33] demonstrated that patients with allocentric neglect had worse functional outcomes in terms of activities of daily living than patients with egocentric neglect. These two points taken together serve to illustrate the severity of this particular type of neglect. Given the separate mechanisms underlying the disorders, traditional interventions aimed at ameliorating egocentric neglect such as cueing and prism therapy are unlikely to be effective and new and innovative rehabilitation strategies may be called for.

In summary, the current study adds to the growing body of evidence viewing neglect as a heterogeneous disorder. There may be a common core spatial deficit, but when it affects egocentric or allocentric space, there are differential and dissociable behavioural profiles as well as differential rates of recovery, with the acute severity of symptoms predicting the persistence of neglect symptoms 6 months later.

## Acknowledgements

We would like to express our sincere thanks and admiration to the late Prof. Glyn W Humphreys, without whose support this study would not have come to pass. We would also like to thank Mr Liam Loftus, who conducted preliminary analyses on an initial subset of this data for his final honours research project, Ms Elitsa Slavkova and Ms Rachel King for help in data collection, and Ms Rachel Teal, stroke research nurse at the John Radcliffe Stroke Unit. This work was supported by the NIHR (grant numbers: RP-DG-0610-10046 & Oxford Cognitive Health CRF); a Stroke Association UK award to N Demeyere (grant number: TSA2015_LECT02); a Sir Henry Wellcome Postdoctoral fellowship to CR Gillebert (grant number: 098771/Z/12/Z); and the Research Foundation Flanders (grant number: G072517N).

## Footnotes

1 All these neuro-anatomical studies were focussed solely on left spatial neglect after unilateral right hemisphere lesions.

2 The allocentric asymmetry cut off represents a conservative approach to reduce potential false positives (see also 17). The 5^th^ centile cut off from the normative sample was 0.

